# Replicative history as a major determinant of epigenetic noise across human tissues

**DOI:** 10.64898/2026.08.19.745715

**Authors:** Alfonso Peñarroya, Juan José Alba-Linares, Raúl F. Pérez, Agustín F. Fernández, Mario F. Fraga, Juan Ramón Tejedor

## Abstract

DNA methylation changes accumulate with age through both regulated and stochastic processes, yet the determinants of epigenetic information loss remain poorly defined. Using genome-wide DNA methylation profiles from 1,531 healthy human samples spanning 14 tissues, we quantified epigenetic noise by Shannon entropy and corrected it for cellular and tissue heterogeneity. Adjusted entropy was consistently low in promoters, first exons and CpG islands, and high in CpG-poor and intergenic regions. Cumulative mitotic history showed a stronger association with epigenetic noise than chronological age, explaining most of its variance particularly within CpG-rich regulatory regions. By contrast, age-related, replication-independent effects predominated outside CpG islands and in low-proliferative tissues such as the brain. Moreover, biological age acceleration was largely attributable to cell division in a tissue-specific manner. Collectively, mitotic history emerges as a major determinant of epigenetic noise accumulation across human tissues, while genomic context modulates regional vulnerability to methylation information loss during aging.

## 1 Introduction

DNA methylation at CpG dinucleotides is a central component of epigenetic regulation, contributing to the maintenance of cellular identity while remaining dynamically modifiable over time[1, 2]. Longitudinal changes in the methylome arise from both regulated processes, including development[3, 4] and environmental responses[5, 6], and stochastic alterations whose origins and functional consequences remain incompletely understood[7, 8]. Collectively, these changes contribute to epigenetic drift[9–12], whereby DNA methylation trajectories progressively diverge among individuals over time[11, 13] under the combined influence of environmental exposures[5] and intrinsic determinants such as genomic context and stochastic errors[14, 15].

Random fluctuations in epigenetic marks are commonly referred to as epigenetic noise[9, 10]. Although subtle at the individual level, the cumulative effects of these stochastic alterations can perturb gene regulatory networks and contribute to age-related functional decline[16–20]. In this context, Shannon entropy provides a useful framework for quantifying stochastic variability in DNA methylation, as it measures the degree of uncertainty or disorder in methylation states using an informational perspective[21]. Within a cell population, higher DNA methylation entropy levels across CpG sites reflects increased heterogeneity and a loss of coherent epigenetic information, reaching its maximum when methylated and unmethylated DNA molecules are present in equal proportions at a given locus. Despite its widespread use as a proxy for epigenetic variability[7, 22], entropy estimates derived from bulk samples are sensitive to cellular composition biases, as shifts in cell-type proportions can alter DNA methylation levels and thereby inflate apparent variability[14, 15]. Consequently, the integration of cell-type deconvolution methods has become essential to disentangle genuine stochastic variation from compositional effects[23].

More broadly, stochastic variability across the lifespan likely reflects compromised fidelity of the molecular mechanisms maintaining DNA methylation patterns. During aging, accumulated damage affects all molecular layers of the cell, including the epigenome[24], driving a progressive loss of epigenetic information causally linked to aging phenotypes[25]. However, variation in methylation entropy is poorly explained by chronological age alone[22], suggesting that environmental exposures, genetic determinants[26], and additional intrinsic processes contribute to its accumulation. In this context, mitotic activity has emerged as a plausible primary driver of stochastic epigenetic change[13], a notion reinforced by the advent of DNA methylation clocks that accurately reconstruct mitotic history[27, 28]. Accordingly, tissues with higher proliferative rates exhibit both increased cancer risk[29] and greater epigenetic variability[30], suggesting a link between replicative history and methylation divergence. However, the quantitative contribution of cumulative cell divisions to epigenetic noise remains insufficiently characterized, particularly across distinct genomic contexts. Recent cross-tissue atlases have reported an age-associated increase in DNA methylation disorder across human tissues, but have not investigated the factors underlying this phenomenon[31].

In this study, we investigated how DNA methylation noise patterns are distributed across the human genome and examined the intrinsic determinants of epigenetic information loss. To address this, we analyzed DNA methylation profiles from thousands of healthy human samples using a unified multi-tissue framework that captures the accumulation of epigenetic variability and links epigenetic noise to replicative history while accounting for cellular heterogeneity.

## 2 Results

### 2.1 Genome-wide landscape of epigenetic noise across human tissues

To establish a comprehensive view of intrinsic epigenetic variability, we first systematically characterized the distribution of DNA methylation noise across multiple human tissues using 1,531 Infinium HumanMethylation450 BeadChip profiles (Fig. 1a, see Materials and Methods). Because these arrays predominantly interrogate single-copy genomic regions, our analyses focused on the unique portion of the human genome. Principal component analysis (PCA, Fig. 1b) and hierarchical clustering (Fig. 1c) revealed distinct tissue-specific DNA methylation landscapes, with peripheral blood and brain showing the greatest separation, consistent with previous studies[32, 33]. Mesenchymal stem cells (MSCs) also displayed marked global hypomethylation relative to differentiated tissues (Fig. 1d), in line with their established epigenetic profile[34, 35].

**Fig. 1.**
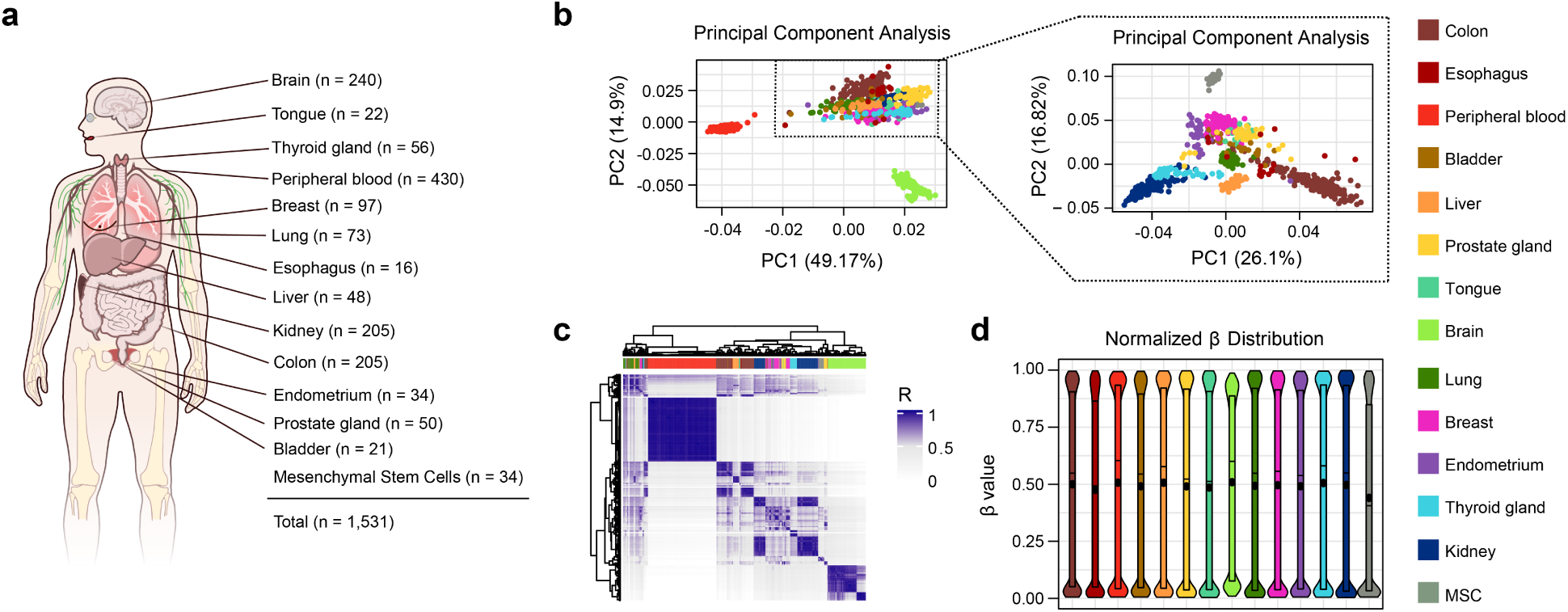
A multi-tissue DNA methylation resource for quantifying epigenetic noise across the human body. **a**, Anatomical overview of the 14 healthy human tissue and cell-type groups analyzed with the Infinium HumanMethylation450 BeadChip platform. Number of samples per tissue are indicated (total *n* = 1,531). **b**, Principal component analysis (PCA) of genome-wide DNA methylation *β*-values. The right panel is a selection of the tissues contained in the boxed area of the original PCA, with the percentage of variance explained shown on each axis. **c**, Heatmap illustrating the unsupervised hierarchical clustering of pairwise sample correlations (Pearson *R*; colour bar) using all the abovementioned tissues. The top annotation bar indicates tissue of origin. **d**, Violin plots indicating the distribution of normalized *β*-values for each tissue. Overlaid box plots show median and interquartile range. The tissue color key applies to panels b–d. Figure 1a has been modified from source. bioart.niaid.nih.gov/bioart/519.

Given the pronounced tissue- and cell-type specificity of DNA methylation patterns across the dataset, we computed Shannon entropy from normalized *β* values and adjusted for tissue and cell-type composition (see Materials and Methods) for each sample. This adjustment enabled quantification of intrinsic stochastic variation in DNA methylation, independent of compositional differences across samples, in accordance with best-practice recommendations[14]. When we examined the genome-wide distribution of DNA methylation noise, we observed a characteristic pattern in which promoter, 5*^′^*UTR, and first exon regions exhibited a valley of adjusted entropy, which increased in gene bodies (avg: +12%, paired t-test *p <* 0.001) and peaked in intergenic regions (avg: +19%, paired t-test *p <* 0.001) (Fig. 2a, top). This trend was consistently found across all analyzed tissues (Fig. 2a, bottom), indicating that it is universal and tissue-independent. Subsequently, we examined DNA methylation noise as a function of CpG density, revealing a pronounced decrease in entropy within CpG islands (CGI; *−*23%, paired t-test *p <* 0.001), which clearly distinguishes them from flanking shores and shelves, as well as Open Sea regions characterized by low CpG density (Fig. 2b, top). This pattern was also conserved across all human tissues (Fig. 2b, bottom).

**Fig. 2.**
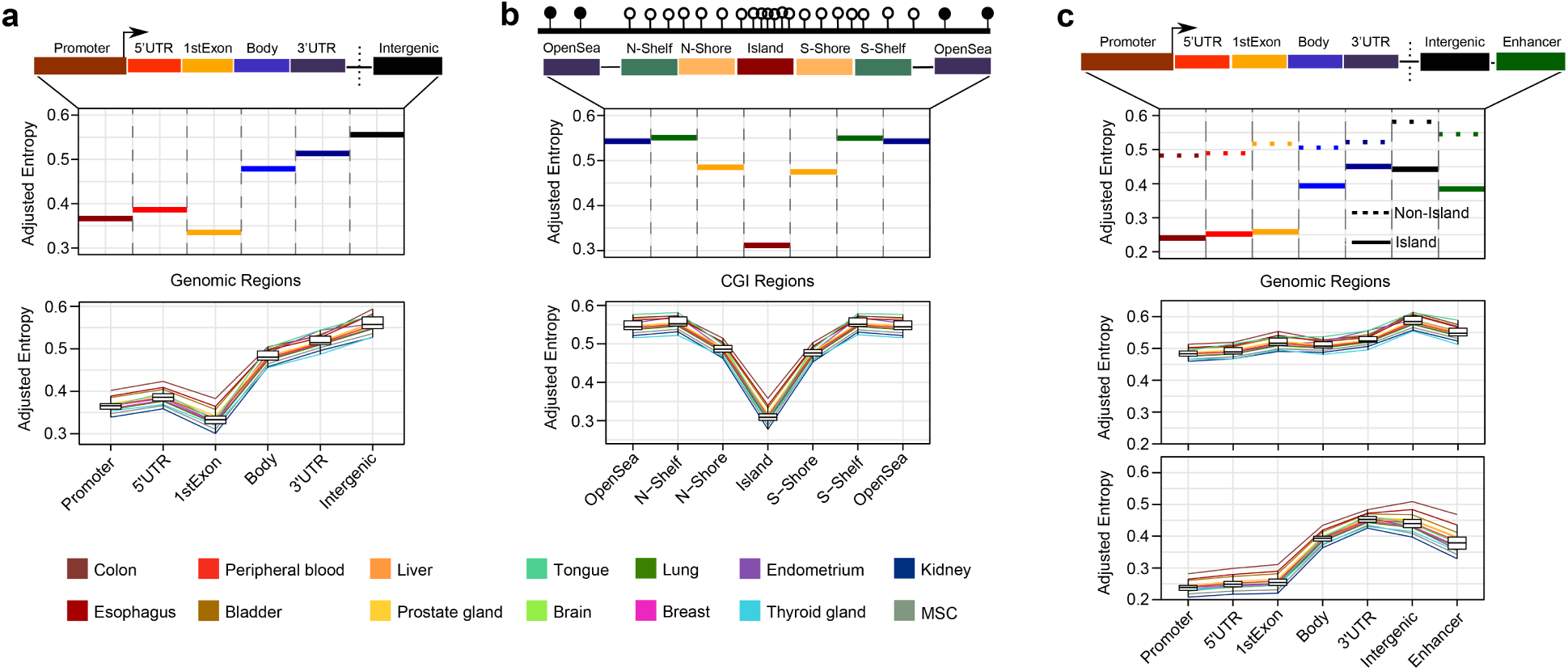
Epigenetic noise is unevenly distributed across genomic contexts and is constrained at CpG islands in a tissue-independent manner. **a**, Top line plot illustrating the adjusted entropy by gene-centric region across all samples (promoter, 5*^′^*UTR, first exon, gene body, 3*^′^*UTR, intergenic). Bottom line plot depicts the per-tissue profiles (one line per tissue; box plots summarize across samples). **b**, Top line plot showing the adjusted entropy across CpG-island (CGI) regions (open sea, N/S shelf, N/S shore, island). Bottom line plot illustrates per-tissue profiles. **c**, Top line plot indicating the adjusted entropy by gene-centric region stratified by CpG-island status (Non-Island, dashed; Island, solid) and including enhancers. Middle and bottom line plots indicate per-tissue profiles of Non-Island and Island CpGs, respectively. Percentage differences quoted in the main text are mean paired differences (paired t-test, *P <* 0.001). Tissue color key is indicated in the legend.

Given that both CpG density and gene context shape DNA methylation noise, we next sought to disentangle the interplay between these sequence-based factors. The entropy valley observed in promoter, 5*^′^*UTR, and first exon regions can be primarily attributed to the presence of CGIs (avg: *−*25%, paired t-test *p <* 0.001), as gene regions lacking them exhibit uniformly higher levels of epigenetic noise (Fig. 2c, top). CGI^+^ promoters displayed low levels of methylation and entropy, whereas their CGI*^−^* counterparts exhibited dual behavior (Supplementary Figure 1). Those with low CpG density presented intermediate levels of methylation that increased epigenetic noise, while those with high CpG density showed low levels of methylation and entropy, and similar patterns were reproduced in 5*^′^*UTR regions (Supplementary Figure 1). Importantly, this pattern was tissue-independent (Fig. 2c, bottom), supporting a central role of CpG islands in constraining entropy across the human genome. Nevertheless, we also identified a contribution of gene context to entropy patterns, as differences between CGI^+^ and CGI*^−^* regions were attenuated in gene bodies (*−*11%, paired t-test *p <* 0.001) and 3*^′^*UTR regions (*−*7%, paired t-test *p <* 0.001) (Fig. 2c, top), demonstrating that gene context also modulates epigenetic noise. Beyond gene-associated regions, CGI^+^ intergenic regions and enhancers exhibited reduced entropy relative to their CGI*^−^* counterparts (*−*14% and *−*16%, both paired t-test *p <* 0.001, respectively), and CGI*^−^* enhancers consistently showed higher entropy levels across all analyzed tissues (ranging from +11% to +21%, Fig. 2c, bottom, Supplementary Figure 1). Overall, our results indicate that CpG-rich regions and promoter-proximal sequences are resistant to the accumulation of DNA methylation noise, a feature that is universal across human tissues.

**Supplementary Figure 1.**
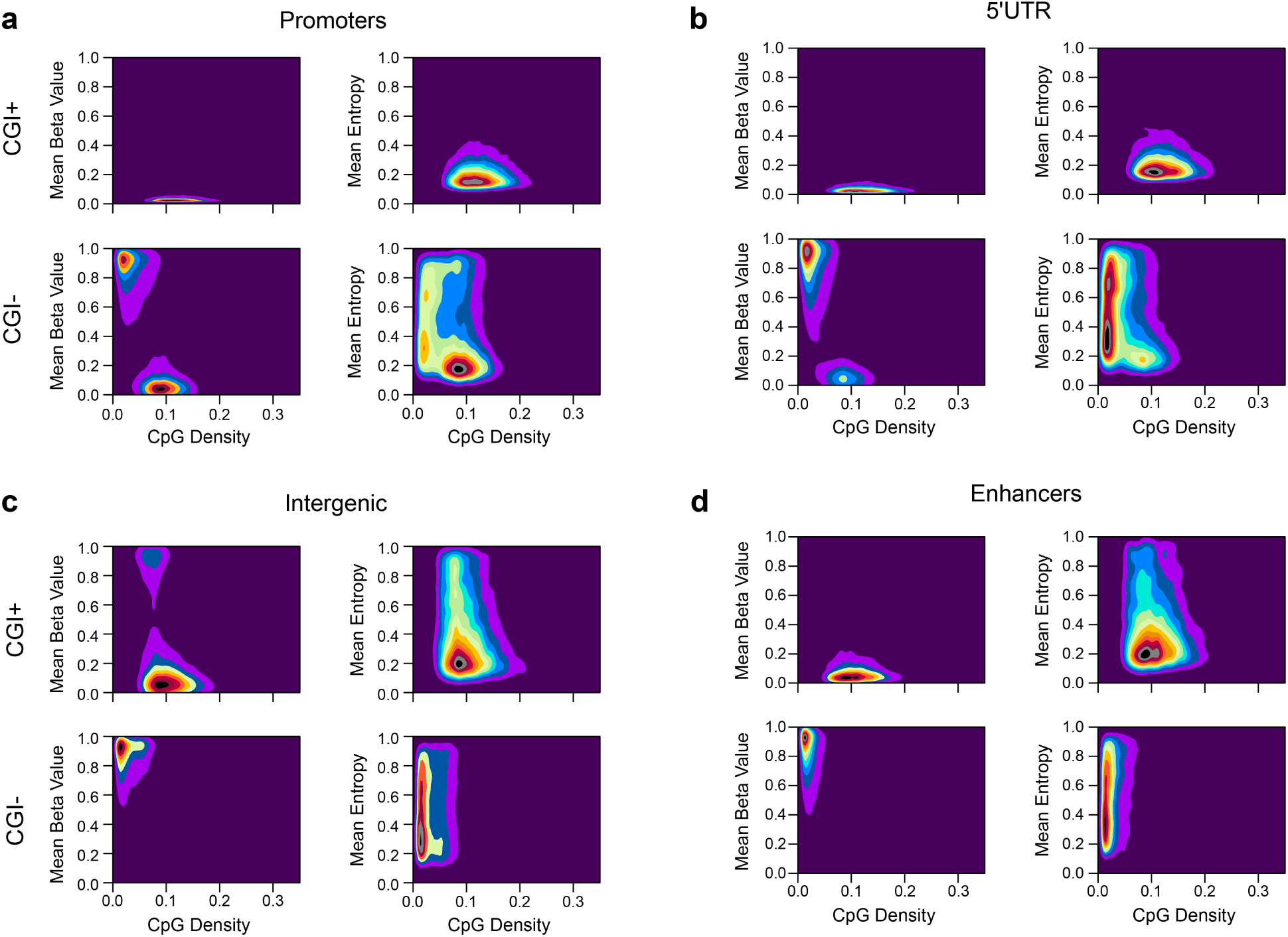
CpG density, rather than CpG-island annotation, defines DNA methylation and entropy behavior across genomic contexts. **a–d**, Two-dimensional density plots indicating mean *β*-value (left) and mean entropy (right) as a function of CpG density, for (**a**) promoters, (**b**) 5*^′^*UTRs, (**c**) intergenic regions and (d) enhancers, each split into CGI^+^ (top) and CGI*^−^* (bottom). Color temperature indicates point density.

### 2.2 Replicative history is a major determinant of epigenetic noise across healthy human tissues

After characterizing the genomic features that shape the baseline distribution of epigenetic noise, we sought to identify the intrinsic determinants of DNA methylation entropy, after accounting for tissue- and cell-type-specific contributions to epigenetic variability. To this end, we evaluated models incorporating biological factors that may contribute to epigenetic noise, including accumulated cell divisions and tissue heterogeneity, as well as global variables capturing stochastic, time-dependent, and environmentally-driven processes, such as chronological age and biological age acceleration (AgeAccel). To emphasize the stochastic, non-deterministic component of aging, we employed first-generation epigenetic clocks across our dataset, including the Horvath and Hannum clocks as measures of biological age, as these clocks are thought to derive much of their predictive accuracy from stochastic age-associated changes in DNA methylation[13]. As expected, these clocks showcased high correlations with chronological age (Supplementary Figure 2a: *r* = 0.7778, *p <* 0.001 for Horvath and *r* = 0.9499, *p <* 0.001 for Hannum, only using peripheral blood samples).

**Supplementary Figure 2.**
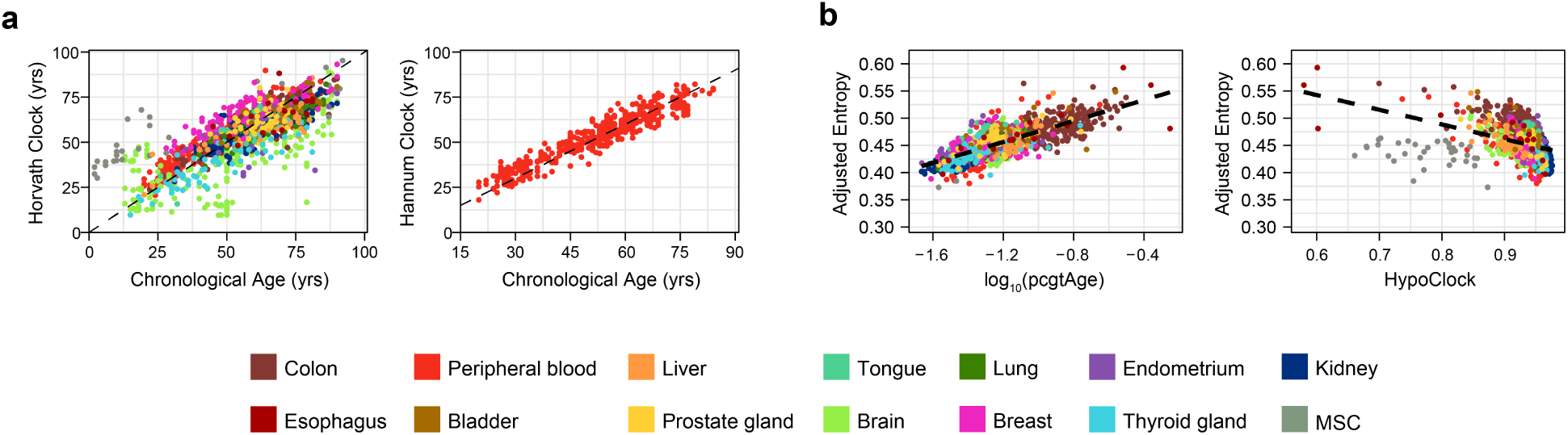
Validation of biological-age and mitotic-clock estimates. **a,** Scatterplots displaying the correlation between chronological age and epigenetic age predicted by the Horvath clock (left, all tissues) and the Hannum clock (right, peripheral blood only). **b,** Scatterplots illustrating the association between adjusted entropy and two additional mitotic clocks: epiTOC (left) and HypoClock (right). Tissue color key is indicated in the legend.

To quantify the contribution of replicative history to epigenetic noise, we employed established mitotic clocks, including epiTOC2[28], to estimate the total number of stem cell divisions accumulated by each sample (TNSC), as well as the intrinsic stem cell division rate per year (irS). In parallel, although entropy estimates were adjusted for tissue- and cell-type composition, tissue complexity and heterogeneity may still holistically influence bulk entropy estimates. To account for this, we defined a composite metric, termed CellScore, that captures the number of distinct cell types within each sample and the similarity of their proportional abundances (see Materials and Methods). Interestingly, chronological age was weakly associated with adjusted entropy (Fig. 3a, *r* = 0.1443, *p <* 0.001), in contrast to mitotic history parameters such as TNSC (Fig. 3b, *r* = 0.7562, *p <* 0.001) and irS (Fig. 3c, *r* = 0.5720, *p <* 0.001). These associations were further supported by additional mitotic clocks (Supplementary Figure 2b), including epiTOC (*r* = 0.7566, *p <* 0.001) and HypoClock (*r* = *−*0.3992, *p <* 0.001).

**Fig. 3.**
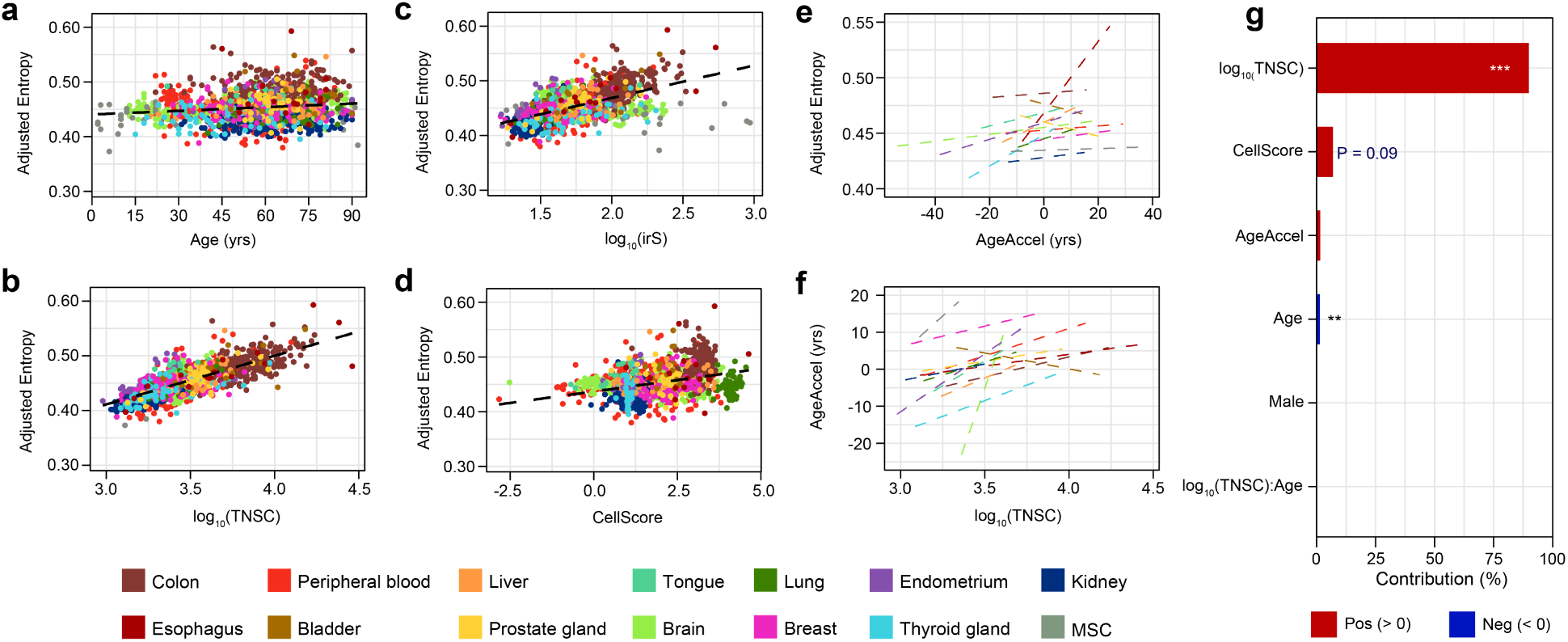
Replicative history is the principal pan-tissue determinant of epigenetic noise. **a–d**, Scatterplots showing sample-level associations between adjusted entropy and (**a**) chronological age, (**b**) cumulative stem-cell divisions (log_10_(TNSC), epiTOC2), (**c**) intrinsic stem-cell division rate (log_10_(irS)) and (**d**) tissue heterogeneity (CellScore). Tissue color key is indicated in the legend. **e**, Line plot indicating the relationship between adjusted entropy and biological age acceleration (AgeAccel), fitted separately within each tissue (one line per tissue). **f**, Line plot depicting the relationship between AgeAccel and log_10_(TNSC) within each tissue. **g**, Barplot illustrating the relative importance of each predictor in a pan-tissue multivariate linear model of adjusted entropy, estimated by variance partitioning (LMG method). Bars show the percentage of explained variance colored by the sign of the association (red, positive; blue, negative). Asterisks and *P* -values denote statistical significance (*^∗∗∗^P <* 0.001, *^∗∗^P <* 0.01).

Moreover, tissue heterogeneity (CellScore) was also positively associated with adjusted entropy levels (Fig. 3d, *r* = 0.3380, *p <* 0.001). Regarding AgeAccel, no robust global association with adjusted entropy was observed across the full sample set (*r* = 0.133), despite highly tissue-specific associations (Fig. 3e; esophagus: *r* = 0.5558, *p <* 0.05, endometrium: *r* = 0.4204, *p <* 0.05, thyroid gland: *r* = 0.4130, *p <* 0.01, liver: *r* = 0.3360, *p <* 0.05). Given that replicative history parameters are also tissue-dependent[29], we hypothesized that AgeAccel dynamics may be driven by the underlying accumulation of cell divisions, a notion we further confirmed (Fig. 3f; most tissues with *r >* 0.30 & *p <* 0.05, including colon, peripheral blood, liver, brain, lung, endometrium, and thyroid gland). In other words, AgeAccel comparisons should be interpreted within a given tissue and are expected to primarily reflect cumulative mitotic history. These associations were further validated using non-adjusted Shannon entropy estimates (Supplementary Figure 3a–d), and were reproduced across individual genomic contexts, including CpG islands (Supplementary Figure 4), confirming that these relationships are not artefacts of the adjustment procedure.

**Supplementary Figure 3.**
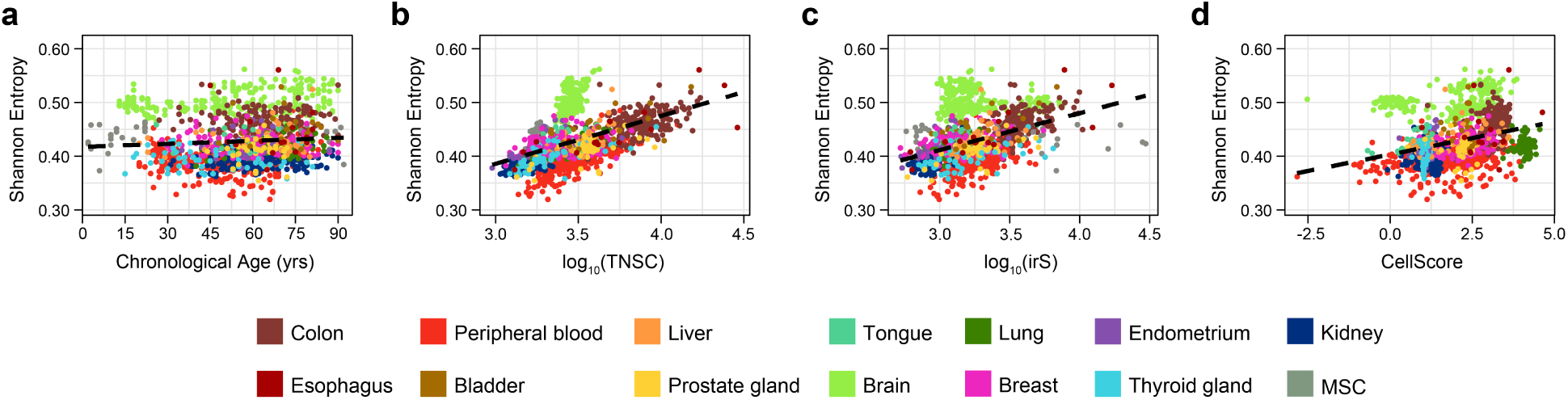
The epigenetic entropy / replicative history association is robust to the entropy definition. **a–d**, Scatterplots illustrating the association between raw (unadjusted) Shannon entropy and (**a**) chronological age, (**b**) log_10_(TNSC), (**c**) log_10_(irS) and (**d**) CellScore. Data demonstrates that the associations in Fig. 3 are not artefacts of the type of adjustment. For a–d, tissue color key is indicated in the legend.

**Supplementary Figure 4.**
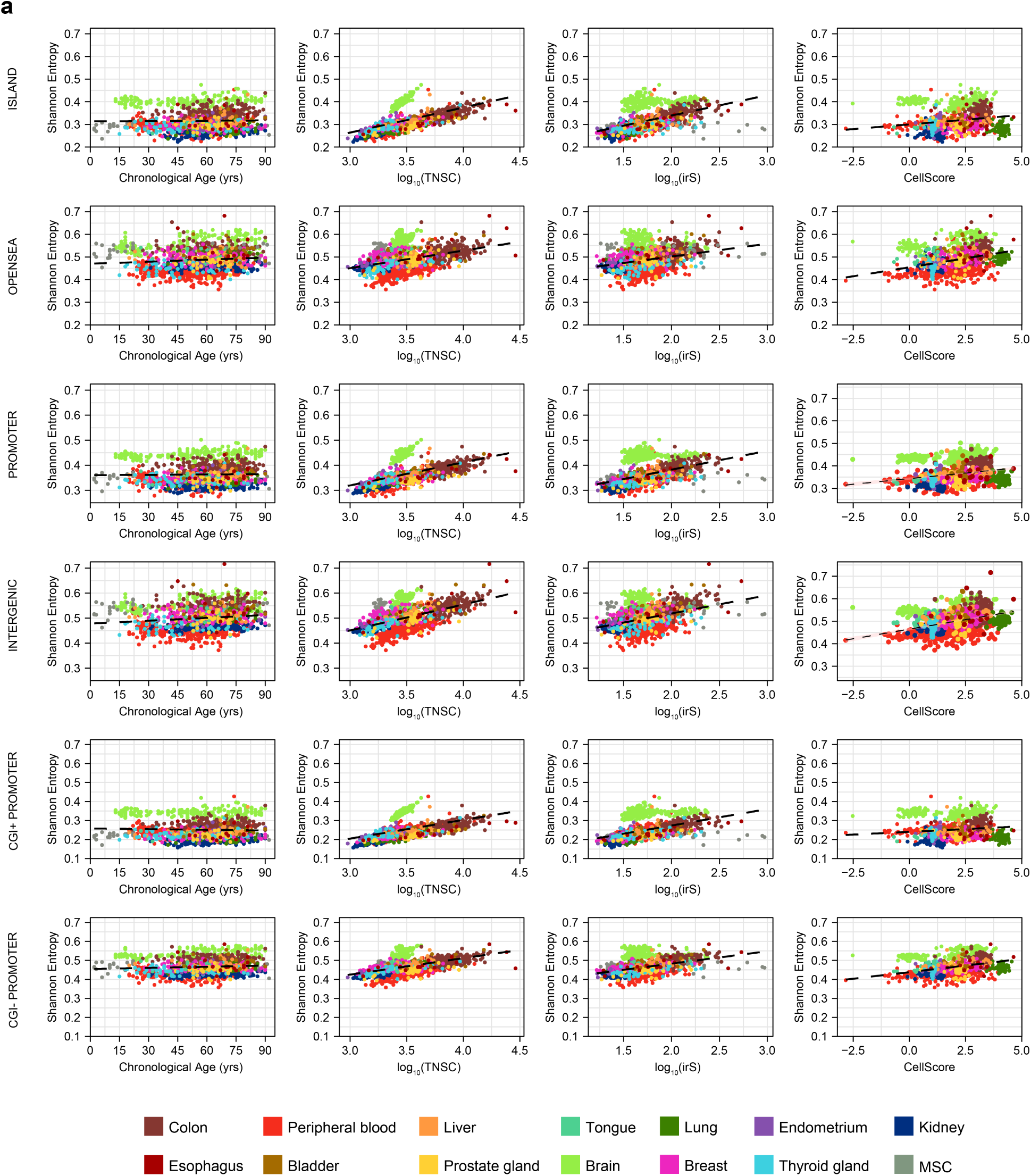
Epigenetic entropy associates with mitotic parameters across all genomic contexts, including CpG islands. **a**, Scatterplots depicting the Shannon entropy versus chronological age, log_10_(TNSC), log_10_(irS) and CellScore, stratified by genomic context across CpG island, open sea, promoter, intergenic, CGI^+^ promoter and CGI*^−^* promoter. Tissue color key is indicated in the legend.

To estimate the relative contribution of all factors studied, we constructed a multi-tissue multivariate linear model predicting adjusted entropy, including TNSC, chronological age, CellScore, AgeAccel, sex, and the interaction between cumulative mitosis and age (TNSC:Age), in light of the well-established age-associated decline in stem cell division rates (Fig. 3g). Interestingly, replicative history accounted for *∼*90.0% of explained variance, followed by CellScore (*∼*7.0%), which showed a near-significant contribution (*p* = 0.09). Although chronological age was significantly associated with adjusted entropy, its effect size was negligible, as were those of AgeAccel and sex. Beyond our pan-tissue model, we additionally constructed 12 tissue-specific multivariate models including the same set of variables whenever possible (Supplementary Figure 5). Across all tissues, DNA methylation entropy was predominantly explained by replicative history, reinforcing our previous observations. In contrast, the contribution of other predictors was less consistent across tissues and generally of limited magnitude. Notably, the brain represented an exception, as the only tissue in which chronological age contributed significantly and to a magnitude comparable to mitotic history (TNSC: 45.3%, age: 29.8%). This finding is particularly relevant given that most brain cell populations are largely post-mitotic in adulthood, suggesting that chronological age may capture replicative-independent sources of epigenetic variability in non-proliferative tissues, potentially related to experience and environment-dependent processes[6].

**Supplementary Figure 5.**
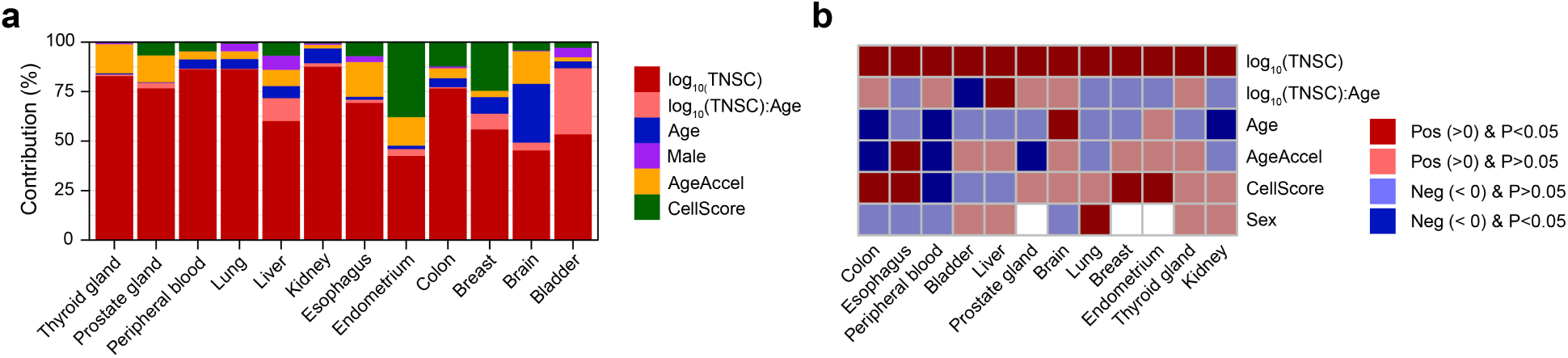
The epigenetic entropy / replicative history association can be decomposed by tissue type. **a**, Stacked bars indicating the relative importance (LMG) of each predictor in tissue-specific multivariate models of adjusted entropy. **b**, Heatmap displaying the direction and significance of each predictor per tissue (red, positive; blue, negative; darker shade, *P <* 0.05).

Altogether, our results indicate that DNA methylation entropy is broadly shaped by the cumulative history of stem cell divisions across human tissues, and that time-dependent epigenetic noise, potentially influenced by environmental factors, is detectable in post-mitotic tissues such as the brain.

### 2.3 Mitosis-induced epigenetic noise particularly targets CpG islands

Having identified replicative history and chronological age as the main determinants of epigenetic information loss, we next investigated how these factors shape the local distribution of DNA methylation noise across the human genome. We first examined adjusted entropy levels in 2-kb windows surrounding CGIs in four representative tissues (Fig. 4a–d): two highly proliferative tissues (peripheral blood and colon), one largely post-mitotic tissue (brain), and mesenchymal stem cells (MSCs). Across all tissues, entropy showed a pronounced decrease within CGIs, consistent with previous observations. We then assessed the local effects of TNSC and chronological age on the distribution of adjusted entropy. In peripheral blood and colon, replicative history was the dominant determinant of adjusted entropy (*>*90%), whereas chronological age contributed negligibly both within CGIs and in flanking regions (Fig. 4a–b). In contrast, in brain tissue, chronological age reached a comparable contribution at distances greater than approximately *±*1 kb from CGI boundaries (from 47% to 68%), while replicative history exerted the strongest influence (99%) within CGIs (Fig. 4c). Finally, MSCs displayed a contribution profile similar to that observed in proliferative differentiated tissues, albeit with greater variability in effect estimates (Fig. 4d).

**Fig. 4.**
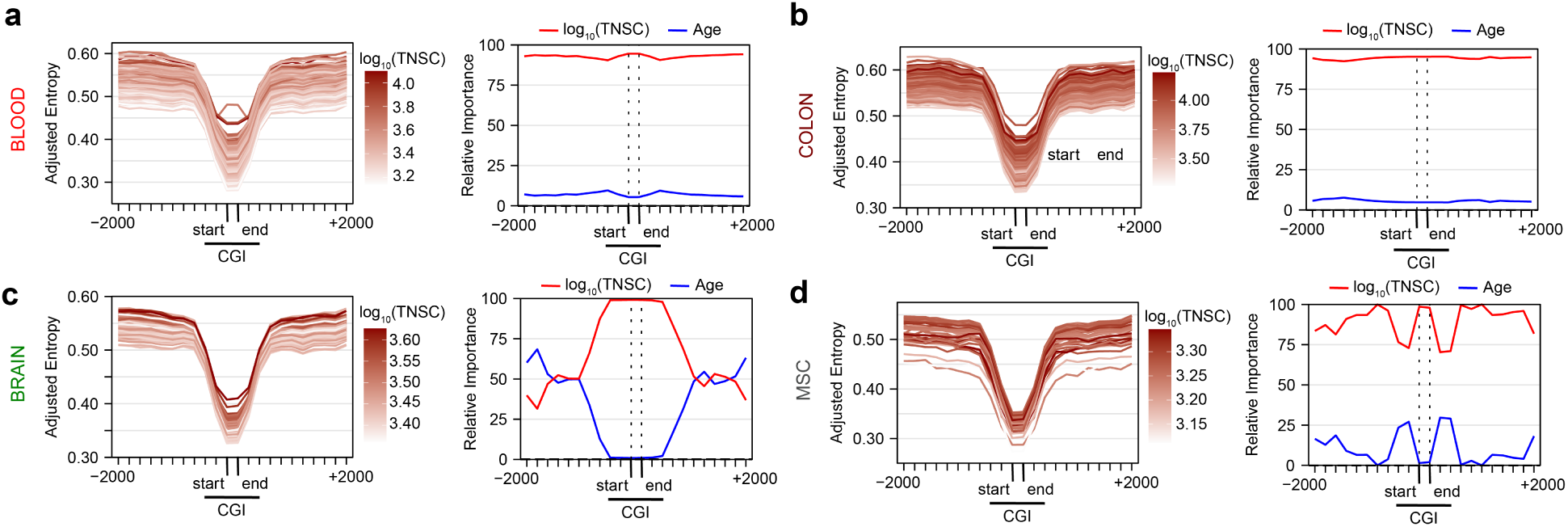
Cumulative cell divisions are the predominant local determinant of adjusted entropy at CpG islands. Line plot showing the local analysis in 2-kb windows flanking CpG islands (CGI; start–end) in (**a**) peripheral blood, (**b**) colon, (**c**) brain and (**d**) mesenchymal stem cells (MSCs). For each tissue, left panel illustrates the adjusted-entropy profiles with samples colored by log_10_(TNSC), and right panel depicts the relative importance (%) of log_10_(TNSC) (red) and chronological age (blue) as determinants of adjusted entropy at each genomic position.

Having established that cell divisions are associated with substantial increases in CGI-targeted epigenetic noise, we next investigated the local contributions of TNSC and chronological age around transcriptional start sites (TSSs) and their flanking regions (*±*2 kb), stratified by the presence or absence of CGIs (Fig. 5a–d). Consistent with our previous observations, adjusted entropy exhibited a pronounced decrease around CGI^+^ TSSs, whereas CGI*^−^* TSSs showed only a modest reduction in entropy immediately downstream of the TSS (blood, brain, MSC: *−*11%, colon: *−*9%). Across the highly proliferative blood and colon tissues, cumulative cell divisions were the primary contributors to increases in adjusted entropy across TSSs and their flanking regions, irrespective of CGI status (CGI^+^: 93% blood, 95% colon, 91% brain, 87% MSC; CGI*^−^*: 91% blood, 95% colon, 87% brain, 94% MSC). Notably, brain tissue and MSCs displayed more variable patterns of relative contribution across flanking regions. Nevertheless, CGI status did not substantially alter these profiles, as replicative history remained the predominant determinant of adjusted entropy across TSS contexts relative to chronological age.

**Fig. 5.**
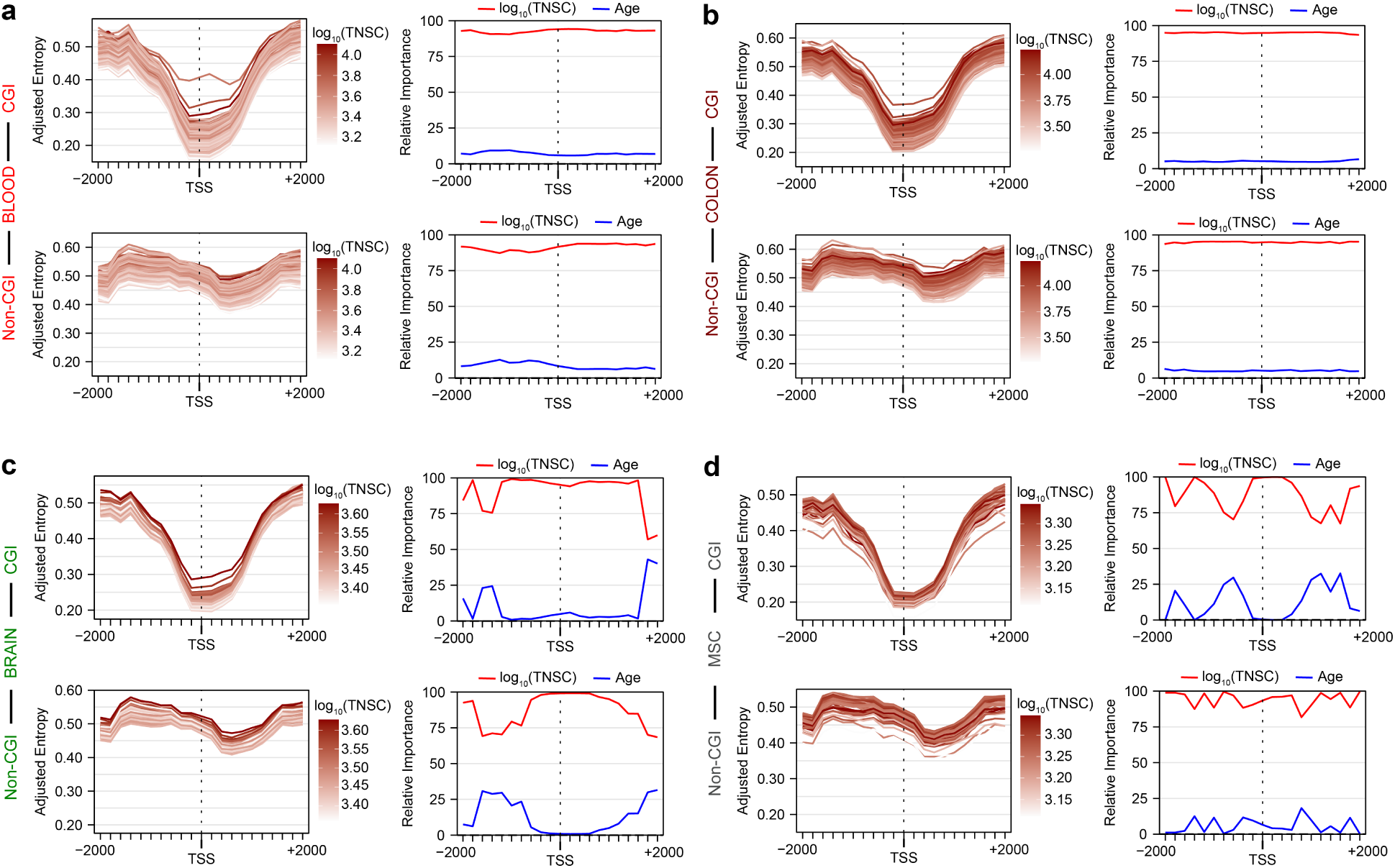
Replicative history dominates adjusted entropy around transcription start sites irrespective of CpG-island status. Line plot indicating the local analysis in 2-kb windows flanking transcription start sites (TSS), stratified into CGI^+^ (top) and CGI*^−^* (bottom) promoters, in (**a**) peripheral blood, (**b**) colon, (**c**) brain and (**d**) MSCs. For each tissue, left panel illustrates the adjusted-entropy profiles with samples colored by log_10_(TNSC), and right panel depicts the relative importance (%) of log_10_(TNSC) (red) and chronological age (blue) as determinants of adjusted entropy at each genomic position.

Together, these findings reveal a context-dependent architecture of epigenetic information loss, whereby replicative history preferentially promotes methylation noise accumulation within CpG islands across all tissues, whereas time-dependent processes contribute predominantly to epigenetic drift outside these regions, particularly in the brain.

## 3 Discussion

The epigenetic changes that occur during aging have been extensively characterized, yet the factors contributing to their emergence remain poorly understood[11, 16, 22, 36, 37]. Here, we examined the determinants of stochastic alterations that erode epigenetic information, collectively referred to as epigenetic noise, across an atlas of 14 healthy human tissues and 1,531 samples. In this context, we focused on several potential contributors, with particular emphasis on cumulative cell divisions, as recent studies have demonstrated that DNA methylation patterns can serve as a record of mitotic history[27, 28, 38]. Importantly, we also developed a framework combining Shannon entropy[39] and cell-type deconvolution analyses[23] to quantify intrinsic DNA methylation disorder independently of age-associated shifts in tissue cell-type composition. This approach enabled us to disentangle the contributions of mitotic and time-dependent, non-replicative factors to epigenetic noise, while controlling for spurious variability introduced by compositional changes in bulk tissues.

Despite the predominantly stochastic nature of epigenetic noise, we observed that entropy patterns are tissue-specific and not uniformly distributed across the human genome. Instead, they are strongly influenced by genomic context and sequence-associated factors, as also observed for mutations[40]. Indeed, we identified a pan-tissue signature of entropy protection at CpG islands and, to a lesser extent, in promoter-proximal regions. Importantly, despite being comparatively protected, we demonstrated that cumulative stem cell divisions are the primary factor explaining entropy accrual during aging across CpG-rich regions in all tissues, including proliferative, stem-cell-like, and post-mitotic contexts. Our results extend the established link between mitotic activity and DNA hypomethylation[41], and further implicate cell divisions in aging-related DNA methylation gains, with CpG islands (which are predominantly unmethylated) emerging as key loci of these changes in human and mouse contexts[42, 43]. Additionally, CpG islands tend to maintain a consistent methylation[42] and transcriptional state[44, 45] across tissues, which may contribute to their preferential association with proliferative history. Finally, our findings have further implications in light of recent evidence showing that DNA hypermethylation leads to stem-cell exhaustion and dysfunction[46], suggesting that replication-driven entropy accrual at CpG islands may promote aging-associated stem-cell phenotypes.

At the genome-wide level, we concluded that replicative history is the primary determinant of epigenetic noise at the pan-tissue level (*∼*90% of explained variance), followed by tissue architecture complexity and heterogeneity, as quantified by our CellScore metric. Interestingly, biological age acceleration derived from first-generation epigenetic clocks, which are relatively enriched for stochastic signals[13], was not informative in multivariate models explaining entropy-based noise once cumulative stem cell divisions were included. Our findings indicate that patterns of biological age acceleration depend on cell division in a tissue-specific manner, with profound implications for the interpretation of epigenetic age acceleration estimates. This is consistent with evidence that accelerated aging in cancer patients is driven by CpG sites in Polycomb-related genes[47], whose methylation levels are profoundly remodeled during cell differentiation processes[48] and replication[27]. However, CpG sites included in the Horvath clock have generally been considered largely independent of mitotic activity[49], based on the weak global correlation between age acceleration and cumulative cell divisions. Of note, as epiTOC2 is derived from methylation gains at PRC2-target CpG islands, part of the island-specific association between TNSC and entropy could in principle reflect a shared genomic substrate. However, the concordant signal obtained with the hypomethylation-based HypoClock (Supplementary Figure 2b), which interrogates late-replicating partially methylated domains, indicates that the mitosis–entropy relationship is not an artefact of any single clock’s design. Our findings suggest that this view may partly reflect the need to account for the tissue-specific nature of biological age acceleration.

Of particular interest is the brain, a post-mitotic tissue in which replicative history was not the absolute driver of entropy levels, in contrast to highly proliferative tissues. In the brain, entropy accrual was significantly associated with time-dependent, replication-independent processes. Given the remarkable epigenetic plasticity of this tissue[50], these findings may reflect the cumulative impact of lifelong environmental exposures, cognitive stimulation, physical activity, memory formation, and other experience-dependent processes that continuously remodel the neuronal epigenome, as has been recently demonstrated in aging contexts[6]. Notably, despite its largely post-mitotic nature, the brain did not exhibit globally lower epigenetic noise than proliferative tissues, but rather reached comparable adjusted-entropy levels through a distinct route. This suggests that mechanistically different sources of epigenetic noise may converge on similar overall levels of methylation disorder by a given chronological age. Under this view, cumulative mitosis and lifelong environmental exposure would represent alternative, partially interchangeable routes to epigenetic information loss, such that tissues with limited proliferative capacity still accrue comparable entropy through replication-independent processes.

Importantly, we observed that these replication-independent contributions to epigenetic noise occurred predominantly outside CpG island regions. Notably, genes located in CpG-island-depleted regions are particularly susceptible to age-related increases in transcriptional noise, a process linked to the loss of cellular identity and chronic inflammation during aging[45]. Consequently, it is plausible that cellular memory and adaptive responses, which increase epigenetic entropy in CpG-island-depleted regions, confer a selective advantage by facilitating environmental adaptation, albeit at the cost of progressively eroding cell-type-specific epigenetic landscapes and fostering cell-to-cell heterogeneity. This increased variability could, in turn, provide a substrate for selection and adaptation at the cell population level, as proposed in models of epigenetic stochasticity in cancer[30, 51]. However, whether this hypothesis extends beyond the brain remains to be determined. Future studies should examine other post-mitotic tissues with substantial epigenetic plasticity, such as adipose and skeletal muscle tissues, as well as stem-cell niches in the bone marrow and colon, which continuously adapt to environmental cues and physiological stressors.

Overall, our work demonstrates that replicative history is the primary intrinsic factor explaining epigenetic entropy dynamics across healthy human tissues, after accounting for cell-type composition biases. Importantly, epigenetic age acceleration can be largely attributed, in a tissue-specific manner, to cumulative cell divisions, highlighting a fundamental link between replication-associated stochastic alterations and biological aging as captured by first-generation epigenetic clocks. CpG islands, which are relatively protected against entropy accumulation, are nevertheless influenced by cell divisions, contributing to age-associated epigenetic drift. Conversely, CpG-island-depleted regions in the brain, a post-mitotic tissue, exhibit substantial replication-independent entropy accrual, potentially driven by environmental and experience-dependent processes.

## 4 Conclusions

This work demonstrates that the cumulative history of stem-cell divisions, rather than chronological age, is the principal intrinsic determinant of DNA methylation noise, and that this replicative contribution is predominant within CpG-rich regulatory regions that are otherwise most protected against entropy accumulation. Replication-independent, time-dependent processes make a comparatively minor genomewide contribution but become dominant outside CpG islands in the post-mitotic brain, where lifelong experience-dependent remodelling likely erodes epigenetic information. As biological age acceleration was largely explained by cumulative mitosis in a tissue-specific manner, our findings argue that first-generation epigenetic-clock signals should be interpreted in the light of tissue proliferative history. Together, these results position cell division as a unifying, quantifiable source of epigenetic information loss during human aging and provide a framework to help disentangle stochastic from programmed methylation changes, with potential implications for future studies focused on the study of aging and cancer.

## 5 Methods

### Description of healthy tissue data sets

Publicly available methylation data (IDAT) from Illumina HumanMethylation450 BeadChip experiments were obtained from the NCBI Gene Expression Omnibus (GEO) and the GDC data portal (The Cancer Genome Atlas, TCGA), including: 430 peripheral blood samples (GSE42861, GSE59065), 240 brain (GSE40360, GSE41826, GSE49393, GSE50798, GSE61107, GSE66351), 205 colon (TCGA-COAD, GSE139404, GSE101764), 205 kidney (TCGA-KIRC, TCGA-KIRP), 97 breast (TCGA-BRCA), 73 lung (TCGA-LUAD, TCGA-LUSC), 56 thyroid gland (TCGA-THCA), 50 prostate gland (TCGA-PRAD), 48 liver (TCGA-LIHC), 34 endometrium (TCGA-UCEC), 34 mesenchymal stem cells (MSC) (GSE52114), 22 tongue (TCGA-HNSC), 21 bladder (TCGA-BLCA) and 16 esophagus (TCGA-ESCA). All considered samples were from healthy tissues annotated with age and sex. All datasets analysed were publicly available and de-identified; informed consent and ethical approval were obtained by the original studies from their respective institutional review boards, so that no additional ethical approval was required for this secondary analysis.

### Methylation array data preprocessing

HumanMethylation450 BeadChip array experiments were analyzed within R statistical software. First, those datasets with raw IDAT files were processed via the minfi package (v.1.44.0)[52], so that methylation data for SNP and sex chromosome probes were accessed to track genetic features and validate self-reported sex, using the getSnpBeta and getSex functions, respectively. Next, array probes were filtered according to the following criteria: (a) detection p-value was *>*0.01 in more than 1% of the samples; (b) those crossreactive or multi-mapping[53, 54]; (c) those located in X/Y chromosomes; and (d) those including SNPs with minor allele frequency *≥* 0.01 at their CpG or single base extension sites (dbSNP v.147). After this, the intensity values from the 390,025 remaining probes that passed all the quality control filters were background corrected using the ssNOOB algorithm[55] (offset=15, dyeCorr=TRUE, dyeMethod=“single”) in minfi. For those samples with data restricted to (un-)methylated probe intensities (datasets: GSE40360, GSE41826, GSE49393 and GSE50798), ssNOOB dye-bias correction was substituted by the nanet approach in the package wateRmelon (v.2.4.0)[56]. Finally, *β*-values were extracted and normalized using the BMIQ method[57] implemented in the package ChAMP (v.2.28.0)[58].

### Probe annotation

The IlluminaHumanMethylation450kanno.ilmn12.hg19 package (v.0.6.0) was used to assign each probe to its CpG Island (CGI) and gene location status. Promoter regions were defined by those CpG annotated as “TSS1500” or “TSS200”. Additionally, transcription start site (TSS) genomic coordinates were extracted from the TxDb.Hsapiens.UCSC.hg19.knownGene package (v.3.2.2) for downstream analyses.

### Epigenetic information loss calculation

Shannon entropy was selected as a theoretical measure to assess the level of biological noise in the methylation layer, and was calculated for each sample as follows[21, 39]:

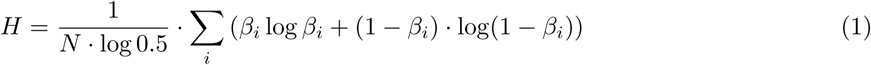

Regional entropy, stratified by gene or CGI status, was computed for each sample using the equation above, restricting the analysis to CpG sites included in the corresponding probe annotations.

According to information theory, when *β*-values are equal to 0 (complete unmethylation) or 1 (complete methylation), the methylation state is uniform across all DNA molecules, and information content is maximal, yielding a Shannon entropy of zero (*H* = 0). In contrast, when the *β*-value is 0.5, predictability of the methylation state within the cell population is minimal, as heterogeneity and thus entropy is maximal (i.e., 50% of DNA molecules are methylated and 50% are unmethylated; *H* = 1).

### Biological age estimation

Horvath and Hannum methylation clocks were used to calculate sample biological age using the methyAge function of the ENmix package (v.1.32.0)[59]. Since methylation clocks tend to underestimate biological age at higher chronological ages[60], biological age acceleration (AgeAccel) was defined as the residuals from a linear regression of Horvath age on chronological age.

### Estimation of cumulative stem cell divisions using epigenetic mitotic clocks

Mitosis levels of the stem cell population, integrating each tissue and individual sample, were inferred using multiple DNA methylation-based clocks. First, the total number of stem cell divisions (TNSC) per sample was estimated using methylation data from the 163 PRC2-associated CpG sites that define the epiTOC2 clock and undergo DNA hypermethylation during aging[28]. Furthermore, the average lifetime intrinsic stem cell division rate (irS) was calculated by dividing TNSC by the chronological age of each sample. In addition, complementary mitotic clocks, such as epiTOC[49], based on 385 age-hypermethylated CpG, and HypoClock[28], based on 678 hypomethylated CpG in partially methylated domains[41], were used to further validate estimates of mitotic parameters.

### Cell-type deconvolution from DNA methylation data

Inferring cell-type composition within each bulk tissue was essential to adjust for potential non-intrinsic variation affecting Shannon entropy levels. Therefore, cell-type proportions were estimated for each sample via the EpiScore package (v.0.9.2)[23] with a weighted robust partial correlation approach that incorporates tissue-specific DNA methylation-based reference matrices. These matrices include gene signatures derived from single-cell RNA-seq atlases, where expression is linked to the methylation state of regulatory regions (promoters and first exons).

In parallel, for solid tissues lacking specific deconvolution references (endometrium, tongue, and thyroid gland), intra-sample heterogeneity was approximated using the EpiDISH package (v.2.4.0)[61] to estimate the fibroblast, epithelial, and immune infiltration proportions. Finally, for peripheral blood, cell-type deconvolution was performed via the EpiDISH suite with the centDHSbloodDMC.m reference dataset.

After cell-type deconvolution, adjusted entropy was defined as a pan-tissue normalized entropy measure that accounts for baseline differences in Shannon entropy across tissues and for variations in cell-type proportions. Accordingly, adjusted entropy values were computed using the removeBatchEffect function from the limma package (v.3.44.3)[62], with per tissue cell-type proportions included as covariates to be adjusted for while preserving variables of interest incorporated in the subsequent models.

### Tissue heterogeneity estimation with CellScore

To quantify tissue variability and complexity from cell-type proportions in a pan-tissue manner, we defined CellScore as the base-10 logarithm of the inverse of a nested ratio of ordered cell-type proportions:

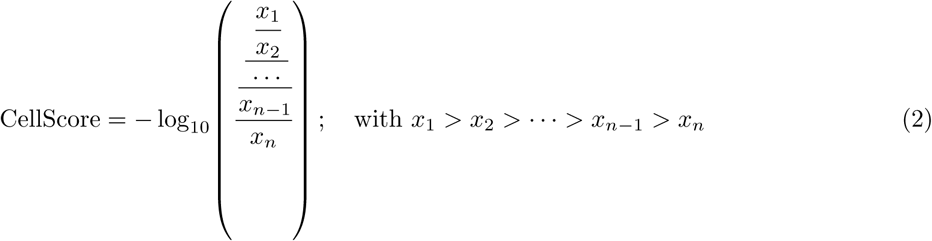

In this formulation, dominance by one or a few cell types, together with a reduced diversity of cell categories, reflects lower tissue heterogeneity and consequently results in lower CellScore values.

### Linear modeling and variable contribution

Multiple linear regression models were constructed in R to assess the relative contribution of distinct factors in predicting adjusted entropy. The following predictors were included to explain epigenetic noise: sex, chronological age, biological age acceleration (AgeAccel), total number of stem cell divisions (TNSC) and the CellScore estimator of tissue heterogeneity.

As mitotic and age-related predictors are partly collinear, the relative importance of each variable in the multivariate models was assessed by partitioning the total explained variance using the Lindeman, Merenda, and Gold (LMG) method implemented via the cal.relimp function from the relaimpo package (v2.2-7)[63], which distributes shared variance across predictors.

### Local-scale epigenetic noise analyses

Adjusted entropy trends were computed for each sample within target regions (CpG islands and transcription start sites) using the normalizeToMatrix function from the EnrichedHeatmap package (v.1.18.2)[64], with the following parameters: extend=2000, w=200, background=NA, smooth=F (for CGI analysis: target_ratio=0.10). Accordingly, epigenetic noise patterns were assessed at the local level by extending each locus into a 2 kb window flanking the region of interest, using 200-bp bins.

### Statistics and Reproducibility

All statistical analyses were performed in R. The relative importance of each predictor was quantified by variance partitioning (LMG method, relaimpo package) in pan-tissue and tissue-specific multiple linear regression models. No statistical method was used to predetermine sample size; sample sizes correspond to all publicly available samples passing quality control for each tissue (*n* = 1,531 in total, with per-tissue numbers indicated in Fig. 1a and the Methods). No samples were excluded after quality control, and all analyses were performed on independent biological samples across the 14 tissue and cell-type groups.

## Data availability

Raw DNA methylation data was obtained from publicly available repositories from the NCBI Gene Expression Omnibus (GEO) and the GDC data portal (The Cancer Genome Atlas, TCGA). Datasets are listed, together with their corresponding NCBI GEO Series (GSE) and TCGA accession codes, in the Methods section (Description of healthy tissue data sets subsection).

## Code availability

No custom software was generated in this study. All analyses were carried out using publicly available, previously published R packages, cited in the Methods and used with the versions and parameters specified therein. Additional analysis scripts are available from the corresponding authors upon reasonable request.

## Acknowledgements

We would like to thank all the members of the Cancer Epigenetics and Nanomedicine laboratory (FINBA-ISPA, IUOPA, CINN-CSIC) for their positive feedback and helpful discussions. We also acknowledge support from the Institute of Oncology of Asturias (IUOPA, supported by Obra Social Cajastur Liberbank, Spain), the Health Research Institute of Asturias (ISPA-FINBA) and Consorcio Centro de Investigación Biomédica en Red (CIBERER-ISCIII).

## Funding

This work was supported by the Spanish Association Against Cancer (PRYGN235109FERN to M.F.F.), the Asturias Government (PCTI) co-funding 2018-2023/FEDER (IDI/2021/000077 and IDI/2024/000744 to M.F.F.), the ISCIII (PI21/01067 and PI24/00641 to M.F.F. and A.F.F. and co-funded by the European Union), the CIBERER, Acciones Cooperativas y Complementarias Intramurales (ACCI20-34-U766 to M.F.F. and ACCI23-13-766 to J.R.T.), ISPA and the Galbán Association (2023-165-GALBAN-TEVAJ to J.R.T.), the eprObes project funded by the European Union through the Horizon Europe Framework Programme (GA 101080219), and ERA-NET TRANSCAN-3 initiative (JTC 2023) (AC24/00163 to M.F.F.). A.P. is supported by the ISCIII (FI19/00085). J.J.A.L. is supported by the AECC foundation. J.R.T. is supported by a Ramon y Cajal fellowship from the Spanish Ministry of Science and Innovation (RYC2021-031799-I).

## Author contributions

A.F.F., M.F.F. and J.R.T. conceived and designed the research. R.F.P., M.F.F. and J.R.T. supervised the study. A.P. and J.J.A.L. designed the computational framework, curated the data, and developed the analytical workflow. R.F.P. and J.R.T. advised on the computational framework. A.P., J.J.A.L., M.F.F. and J.R.T. wrote the manuscript. All authors contributed to the manuscript revision.

## Competing interests

The authors declare no competing interests.

## Notes

### Competing Interest Statement

The authors have declared no competing interest.

