## Supplementary figures and images for "Replicative history as a major determinant of epigenetic noise across human tissues"

### Supplementary Figure 1

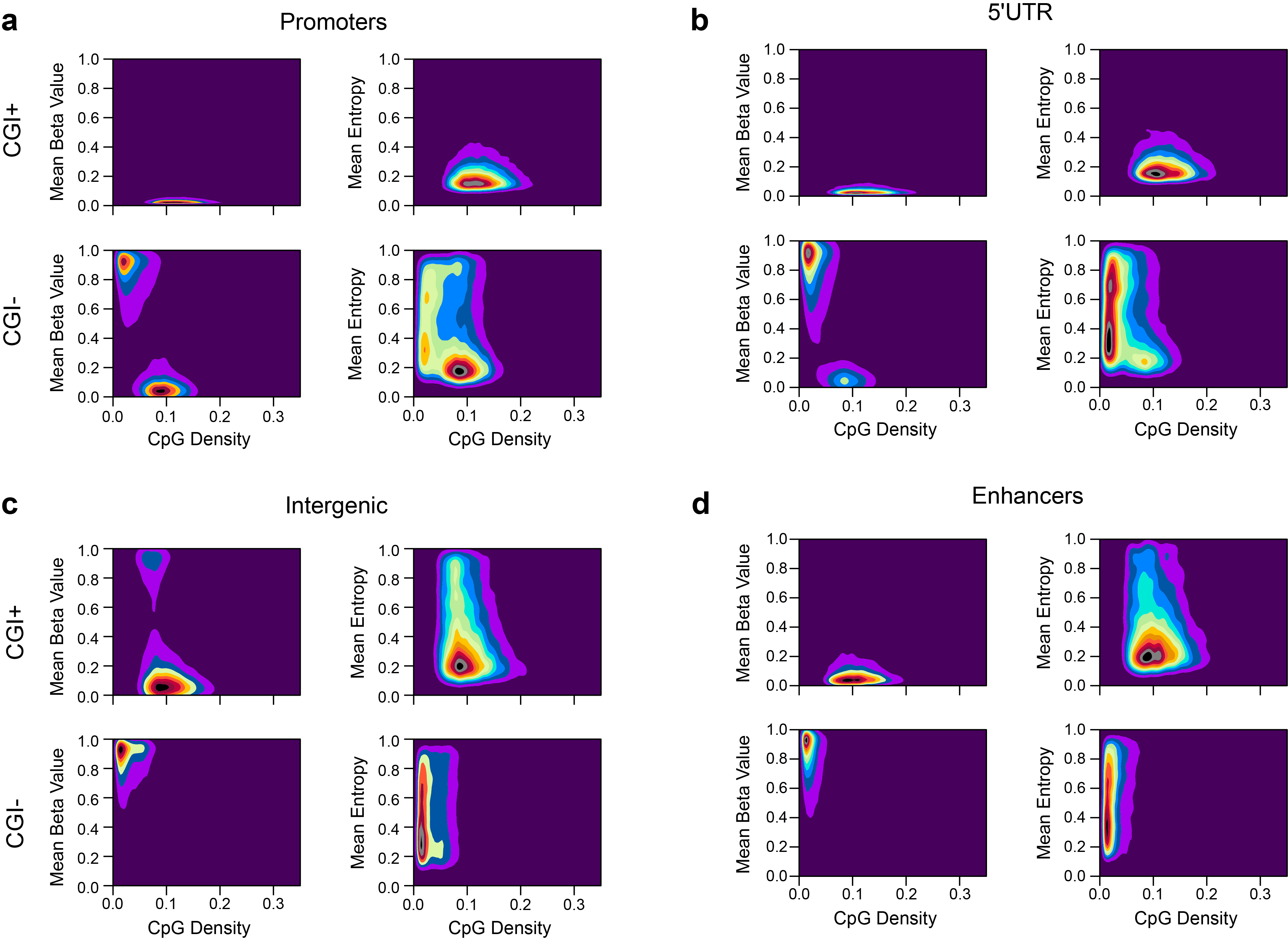

### Supplementary Figure 2

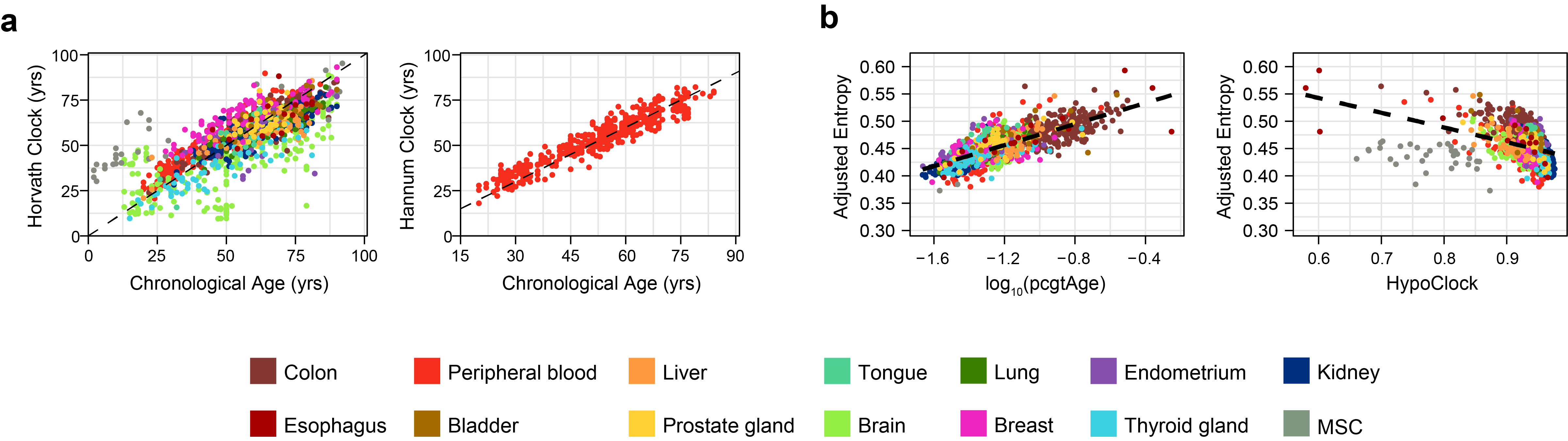

### Supplementary Figure 3

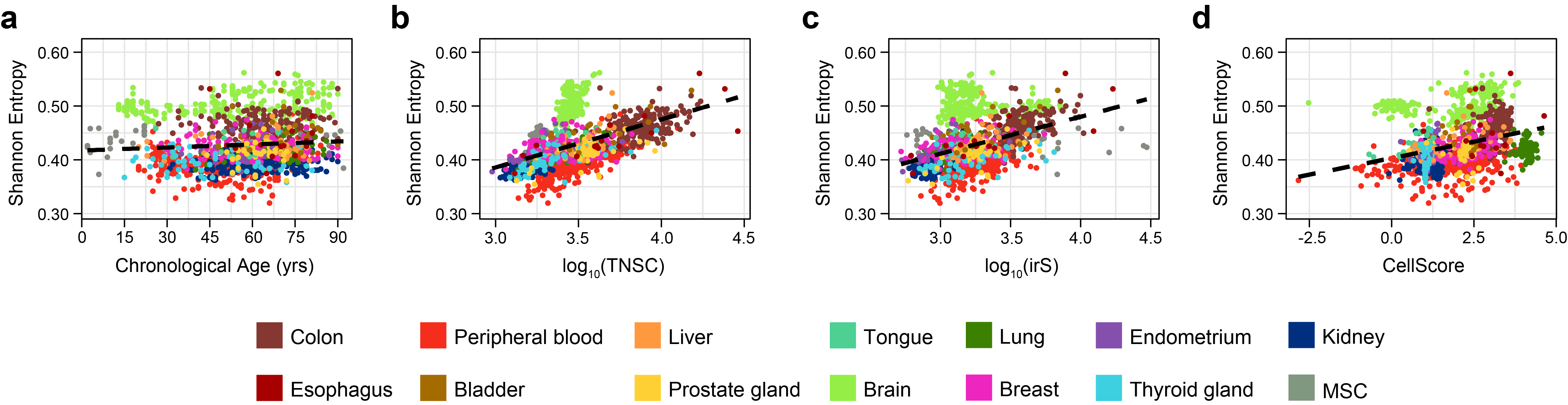

### Supplementary Figure 4

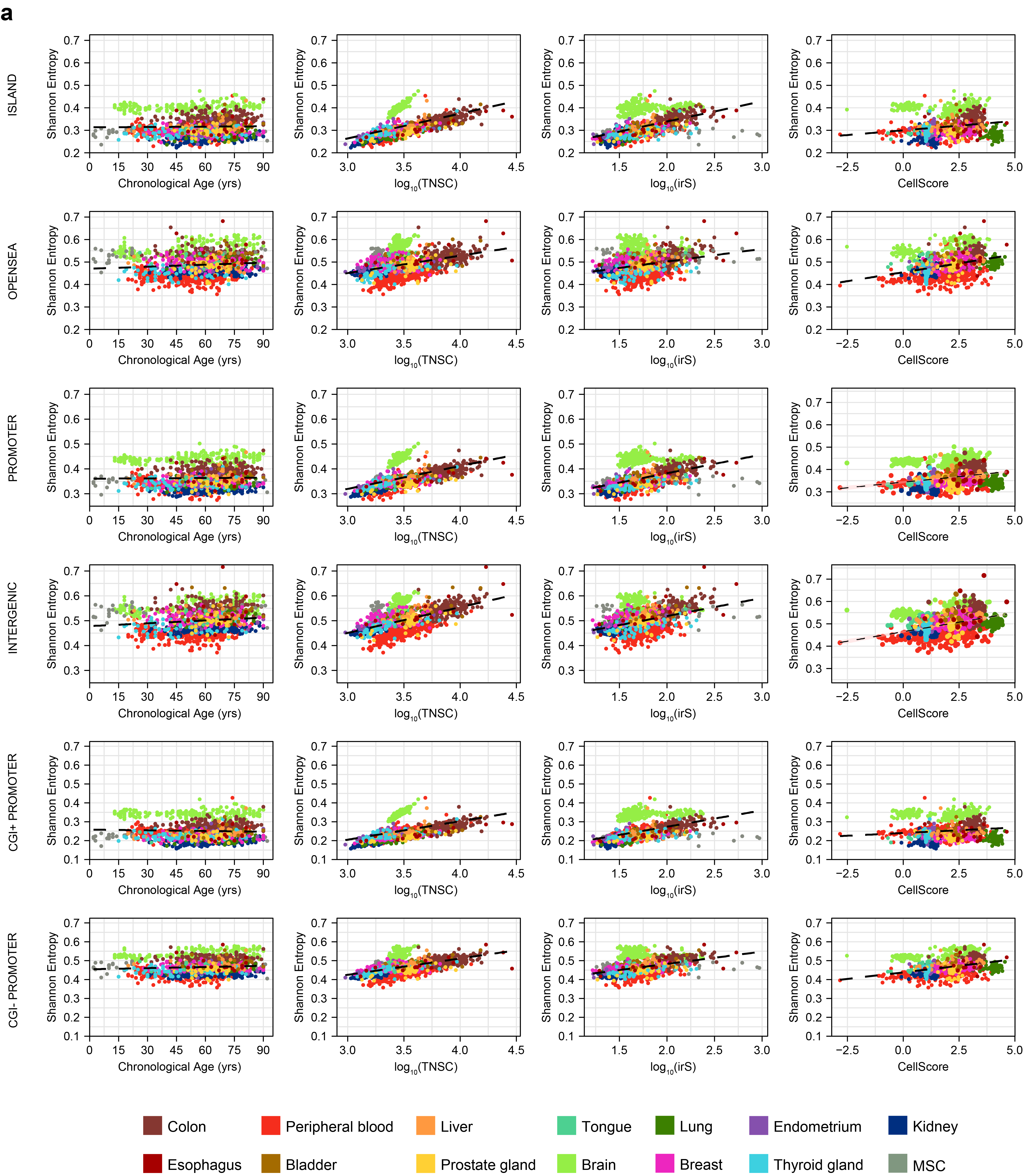

### Supplementary Figure 5

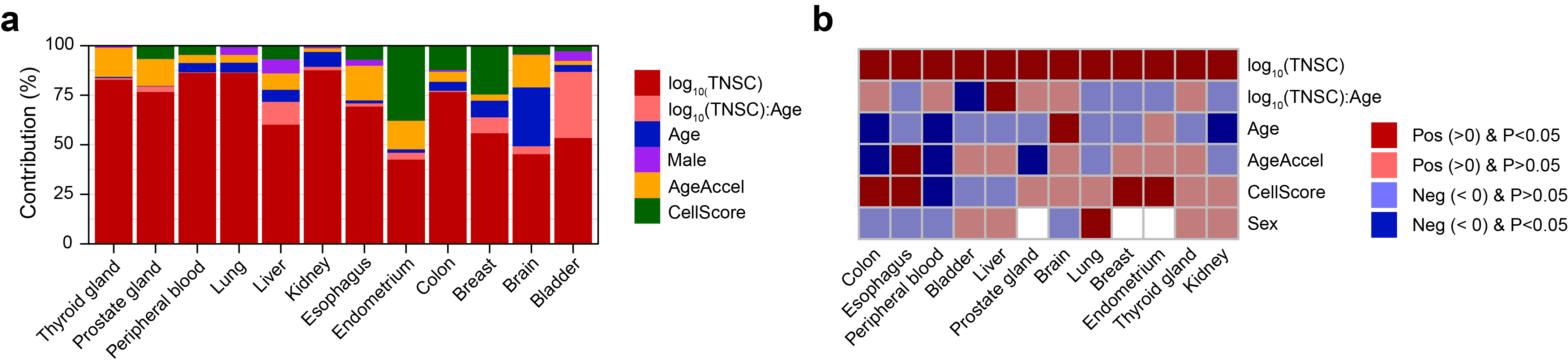
